# The effects of cannabidiol on cue- and stress-induced reinstatement of cocaine seeking behavior in mice are reverted by the CB1 receptor antagonist AM4113

**DOI:** 10.1101/2020.01.23.916601

**Authors:** Miguel Ángel Luján, Laia Alegre-Zurano, Ana Martín-Sánchez, Olga Valverde

**Author notes:** Author for correspondence: Olga Valverde, MD PhD, Neurobiology of Behaviour Research Group (GReNeC - NeuroBio), Department of Experimental and Health Sciences, Universitat Pompeu Fabra, Dr. Aiguader 88; Barcelona 08003.

## Abstract

Cocaine addiction is a brain disorder characterized by the consumption of the drug despite harmful consequences, the loss of control over drug intake and increased risk of relapse. Albeit prolonged research efforts, there is no available medication approved for the treatment of cocaine addiction. In the last decade, cannabinoid-based compounds have drawn increased interest for its potential therapeutic applications in various psychiatric conditions. Cannabidiol, a non-psychotomimetic constituent of the *C. sativa* plant, shows promising results in rodent models of anxiety, schizophrenia, depression and drug addiction. However, the specific effects and mechanisms of action of cannabidiol in a rodent model of extinction-based abstinence and drug seeking relapse remain unclear. Here, we administered cannabidiol (20 mg/kg) to male CD-1 mice trained to self-administer cocaine (0.75 mg/kg/inf) during extinction training (8–12 days). Then, we evaluated the reinstatement of cocaine seeking induced by cues, stress and drug priming. To ascertain the participation of CB1 receptors in these behavioral responses, we systemically administered the neutral cannabinoid antagonist AM4113 (5 mg/kg) before each reinstatement session. The results document that cannabidiol (20 mg/kg) does not modulate extinction training but attenuates ‘extinction burst’ responding after one cannabidiol injection. Furthermore, cannabidiol specifically blocked the reinstatement of cocaine seeking triggered by a cue presentation, an effect prevented by AM4113 (5 mg/kg). Unexpectedly, cannabidiol facilitated stress-induced reinstatement of cocaine seeking behavior, also by a CB1-dependent mechanism. Finally, cannabidiol did not affect cocaine-primed (10 mg/kg) precipitation of cocaine seeking. Our results reveal a series of complex changes induced by cannabidiol treatment with opposite implications for the reinstatement of cocaine seeking behavior that may limit therapeutic opportunities. The activity of CB1 receptors seems to play a crucial role in the expression of cannabidiol-induced neuroplasticity underlying both the desirable and undesirable reinstatement effects here detailed.

## INTRODUCTION

Drug addiction is a chronic and relapsing disease characterized by the compulsive taking and pursue of the drug, despite its harmful consequences (Koob and Volkow, 2016). Cocaine is the most consumed psychostimulant worldwide and has shown increased markers of prevalence in the last years (European Monitoring Centre for Drugs and Drug, 2019; United Nations, 2019). Despite extensive research on the neurobiological bases of the disease, no approved medication is yet available (Kampman, 2019).

In the recent years, there has been resurged interest in the potential therapeutic benefits and applications of cannabinoids, or cannabis-based products (Fischer et al., 2015). This situation has been further accelerated in the current context of the proliferation of medicinal and recreational cannabis legalization in several countries (Ware, 2018). Nonetheless, prescription of *Cannabis sativa* for medical use faces major challenges (Wilkinson et al., 2016). However, *Cannabis sativa* contains other 120 phytocannabinoid compounds, each one with particular pharmacological characteristics and mechanisms of action (ElSohly and Gul, 2015). Recent advances in the experimental assessment of isolated phytocannabinoids have spurred the search for those constituents that may offer improved opportunities and be more readily amenable for use as therapeutics than Δ^9^-tetrahydrocannabinol (THC) (Chye et al., 2019).

Cannabidiol (CBD), the most abundant non-psychotropic constituent of the *Cannabis sativa* plant, has drawn the attention of scientific community due to its putative desirable effects for the treatment of various psychiatric diseases (Devinsky et al., 2014), including drug addiction (Zlebnik and Cheer, 2016). CBD has been shown able to reduce the behavioural and molecular manifestations of maladaptive neuroplasticity underlying drug addiction (Hurd et al., 2019). However, available evidence supporting a protective role of CBD in pre-clinical models of volitional drug seeking, and its mode of action, is still scarce. So far, two studies have suggested that CBD administration can reduce cocaine intake in mice (Luján et al., 2018) and rats (Galaj et al., 2019). A single report by Gonzalez-Cuevas et al. (2018) supported the treatment potential of CBD in reducing cue- and stress-induced retrieval of cocaine seeking behavior in rats. In contrast, it was also reported that CBD had no effect on cocaine seeking behavior (Mahmud et al., 2017). Inquiries into the responsible mechanisms of action highlight the relevance of CBD’s pro-neurogenic effects (Luján et al., 2019), as well as cannabinoid receptor 2 (CB2), 5-hydroxytryptamine 1A (5-HT_1A_) and transient receptor potential vanilloid 1 (TRPV1) receptor activity (Galaj et al., 2019) to reduce cocaine intake. Until date, no studies have evaluated CBD-induced pharmacological effects underlying its influence on the reinstatement of cocaine seeking behavior (Calpe-López et al., 2019). Pharmacological studies aimed to decipher CBD’s mechanisms of action on the reduction of cocaine seeking reinstatement are crucially needed. This is because reinstatement to drug seeking after extinction-based abstinence in rodents resembles the clinical setting in which most treatment entries occurs (Epstein et al., 2006). Exploring the mechanisms of action by which CBD exerts its protective effects in a rodent model of cocaine relapse would allow us to gain critical insights into the therapeutic strengths, and limitations, of this phytocannabinoid.

CBD is thought to interact with a several cellular and molecular targets (Campos et al., 2017). Its main molecular effects within the central nervous system are comprehended by the agonism of 5-HT_1A_, TRPV1, G-protein receptor 55 (GPR55), peroxisome proliferator-activated gamma (PPARγ) receptors and the antagonism of adenosine reuptake (Turner et al., 2017). Despite initial controversy about its endocannabinoid targets (Zlebnik and Cheer, 2016), recent evidence also supports CBD as a negative allosteric modulator of CB1 and CB2 receptors at physiologically relevant concentrations (Laprairie et al., 2015; Martínez-Pinilla et al., 2017; McPartland et al., 2015; Navarro et al., 2018; Tham et al., 2019). In addition, CBD reduces anandamide (AEA) metabolism by inhibiting fatty acid amid hydrolase (FAAH) enzymatic activity (De Petrocellis et al., 2011). Among these molecular targets, CB1 receptors are of special interest to the motivational effects of CBD. CB1 receptors are uniquely positioned to serve as fine tuners of the mesolimbic dopaminergic activity (Oleson et al., 2012; Wenzel and Cheer, 2018) underlying cocaine-related behaviors. Therefore, compounds that modify CB1 receptor activity are known to modulate reinstatement of cocaine-seeking behavior (De Vries et al., 2001; De Vries and Schoffelmeer, 2005).

In the present study, we evaluate the pharmacological effects of an extinction-targeted CBD treatment on the reinstatement of cocaine voluntary seeking in a mouse model of intravenous self-administration. Moreover, we assess the effects of CBD on cue-, stress- and drug-induced reinstatement of cocaine seeking behavior. To explore a possible CB1 receptor mechanism underlying the expression of CBD effects, we administer the antagonist AM4113 in CBD- and vehicle-treated mice undergoing cocaine-seeking reinstatement.

## METHODS

### Animals

All animal care and experimental protocols were approved by the UPF/PRBB Animal Ethics Committee, in line with European Community Council guidelines (2016/63/EU). Male CD-1 mice (PND 41-44) were purchased from Charles River (Barcelona, Spain). All efforts were made to minimize animal suffering and to reduce the number of animals used. Animal studies are reported in compliance with the ARRIVE guidelines (Sert et al., 2019). Animals were maintained in a 12h light–dark cycle, in stable conditions of temperature (22°C), with food and water *ad libitum.* The CD-1 mouse strain was selected for its optimal sensitivity to the reinforcing effects of cocaine (McKerchar et al., 2005) and its fast recovery from catheterization surgery. A total sample of 68 mice was used for the self-administration studies.

### Cocaine operant self-administration

#### Surgical procedures

The self-administration procedure here employed was conducted as previously described in Tourino et al. (2012) and Luján et al. (2018; 2019). Surgical implantation of the catheter into the jugular vein was performed following anaesthetization with a mixture of Ketamine hydrochloride (100 mg/kg; Imalgène1000, Lyon, France) and Xylazine hydrochloride (20 mg/kg; Sigma Chemical Co., Madrid, Spain). The anesthetics solution was injected in a volume of 0.15 mL/10 g body weight, i.p. Briefly, a 6 cm length of silastic tubing (0.3 mm inner diameter, 0.6 mm outer diameter) (silastic, Dow Corning, Houdeng-Goegnies, Belgium) was fitted to a 22-gauge steel cannula (Semat, Herts, England) that was bent at a right angle and then embedded in a cement disk (Dentalon Plus, Heraeus Kulzer, Germany) with an underlying nylon mesh. The catheter tubing was inserted 1.3 cm into the right jugular vein and anchored with a suture. The remaining tubing ran subcutaneously to the cannula, which exited at the mid-scapular region. All incisions were sutured and coated with antibiotic ointment (Bactroban, Glaxo-SmithKline, Spain). After surgery, mice were housed individually and allowed to recover for at least 3 days. During recovery, mice were treated daily with an analgesic (meloxicam 0.5 mg/kg, injected in a volume of 0.1 mL/10 g, i.p.) and an antibiotic solution (enrofloxacin 7.5 mg/kg, injected in a volume of 0.03 mL/10 g, i.p.). The home cages were placed upon thermal blankets to avoid post-anesthesia hypothermia.

#### Acquisition of operant taking behavior

Active and inactive nosepoke holes were assigned randomly to each animal. Cocaine (0.75 mg/kg) was delivered in a 20 μl injection over 2s via a syringe mounted on a microinfusion pump (PHM-100A, Med-Associates, Georgia, VT, USA) connected to the mouse’s intravenous catheter. All FR1 sessions started with a cocaine priming infusion. When mice responded on the active hole, the stimulus lights lit up for 4s. Each infusion was followed by a 15s time-out period in which active nosepokes had no consequences. Mice were considered to have acquired stable self-administration behavior when the following criteria were met on 2 consecutive FR1 sessions: a) 80% stability in reinforcements; b) ≥ 75% of responses were received at the active hole; and c) a minimum of 8 responses on the active hole. Overall FR1 phase operant responding was quantified as the mean number of responses in the active hole given after having reached acquisition criteria.

#### Extinction of cocaine operant responding and CBD treatment

After training, animals that reached acquisition criteria were moved to the extinction phase 48 h after the last FR1 session. Extinction sessions (2 h) were conducted once a day and responses on the active hole produced neither a cocaine infusion nor stimulus light presentation. Before the first session, animals were randomly assigned to the CBD or vehicle group. CBD (20 mg/kg, i.p.), or vehicle, were systemically administered immediately before the start of each self-administration session. Animals underwent a minimum of 8, and a maximum of 12 extinction sessions and were required to reach extinction criteria to progress to the reinstatement tests. A subject was considered to have extinguished cocaine seeking behavior when performing less than 8 active nosepokes or 40% of the overall FR1 phase responding during two consecutive sessions.

#### Reinstatement of cocaine seeking behavior and CB1 receptor blockade

24 h after extinguishing cocaine seeking behavior, CBD- and vehicle-treated mice were randomly assigned to one out of three reinstatement conditions. Reinstatement test sessions (2 h) were conducted as reported in Soria et al. (2008). Cue-induced reinstatement was precipitated by presenting the cocaine-associated cue light after each active nosepoke. A time-out period of 15 s for light presentation was stablished. Additionally, the sessions started with a non-contingent presentation of the light cue (5 s). For the drug-induced reinstatement sessions, animals were confined to operant chambers immediately after receiving an i.p. cocaine injection (10 mg/kg). Active nosepokes had no consequences whatsoever. Finally, stress-induced reinstatement of cocaine seeking behavior was accomplished by delivering a 2 s footshock (0.21 mA) every minute during a 5 min period (total footshocks: 5). As in the case of cocaine-induced reinstatement, no cue light or cocaine were presented after nosepoking.

Each animal received two reinstatement sessions separated of 48 h. To minimize the number of animals used in the study, the administration of the CB1 antagonist AM4113 (5 mg/kg, i.p.) followed a within-subject design. Each subject performed one reinstatement session after vehicle injection and the other reinstatement session after AM4113 administration. The order in which vehicle and AM4113 were administered was counterbalanced between subjects. AM4113 treatment conditions were the same among the three different sets of animals undergoing cue-, drug- or stress-induced reinstatement sessions.

### Materials

#### Drugs

Cocaine HCl (0.75 mg/kg, 10 mg/kg; Alcaliber S.A., Madrid, Spain) was dissolved in 0.9% NaCl. CBD (20 mg/kg) was provided by courtesy of Phytoplant Research S.L., (Córdoba, Spain) and freshly prepared in a 2% Tween80-containing 0.9% NaCl solution. CBD doses were selected based on previous studies from our laboratory showing an attenuating effect on cocaine taking behavior (Luján et al., 2019; 2018), and were within the range of doses used in previous studies (Fogaça et al., 2018; Ren et al., 2009). AM4113 (5 mg/kg) was purchased from Axon Medchem L.L.C. (Reston, VA, USA) and dissolved in a 0.9% NaCl solution containing 5% cremophor. To avoid an unspecific effect of CB1 receptor blockade on cocaine seeking behavior (De Vries et al., 2001), an ineffective dose of 5 mg/kg (He et al., 2018) was chosen.

#### Apparatus

Self-administration training and testing occurred in operant chambers (Model ENV-307A-CT, MED Associates, Inc., Georgia, VT, USA) equipped with two presence-detecting nosepoke holes. Chambers were made of aluminum and clear acrylic, had grid floors connected to an electrical shocker (ENV-414, MED Associates, Inc., Georgia, VT, USA), and were housed in sound- and light-attenuated boxes equipped with fans to provide ventilation and ambient noise. Two cue lights per nosepoke were used, one located above and the other inside the nosepoke hole.

### Data acquisition and analysis

The data and statistical analysis in this study comply with the recommendations on experimental design and analysis in pharmacology (Curtis et al., 2015). Data were expressed as mean ± SEM. Animals were randomly assigned to an experimental group. During the behavioural manipulations and data interpretation, researchers were blind to the treatment that each animal had received. The exact group size for the individual experiments is shown in the corresponding figure legends. We analyzed the results of single factor, parametric measures with unpaired Student’s *t* tests. Parametric measures resulting from the combination of two factors were analyzed with 2-way ANOVAs. When an experimental condition followed a within-subject design (e.g. *session*, *AM4113 treatment*) a two-way ANOVA with repeated measures was calculated. The α-level of statistical significance was set at *p* < 0.05. When required, ANOVAs were followed by Bonferroni’s post-hoc tests (GraphPad Prism 7, La Jolla, CA, USA). This was done only in the case that the F value reached the level of significance, dependent measures followed a normal distribution (Shapiro-Will’s test), and no significant variance in homogeneity was observed (Bartlett’s test).

## RESULTS

### CBD treatment effects on the extinction of cocaine seeking operant behavior

After being trained to nosepoke for cocaine (0.75 mg/kg/inf) for 10 days (Figure 1a), animals were submitted to an extinction protocol for 8-12 days and were repeatedly treated with CBD (20 mg/kg) immediately before being placed in the operant chambers. A two-way, repeated measures ANOVA analysis of nosepoke responding yielded a significative contribution of the factor *day* (F_9,479_ = 38.11; *p* < 0.001), but not *CBD treatment* (F_1,65_= 0.29; *p >* 0.59). However, a significant interaction between both factors was found (F_9,479_ = 2.45; *p* < 0.01), meaning that CBD exerted day-dependent behavioral effects (Figure 1b). This result was followed by Bonferroni’s multiple comparisons tests. Results indicate that the time-dependent modulation of extinction learning was due to a significant reduction of cocaine seeking in CBD-treated mice on the first day of this experimental phase (Bonferroni; *p* < 0.05). A closer examination to the CBD effect on the first day of extinction is presented in Figures 1c and 1d. Student’s *t* test revealed that after a single CBD (20 mg/kg) injection, cocaine seeking was attenuated in the first extinction session (unpaired *t* test: *t*_66_ = 2.01; *p* < 0.05) (Figure 1c). A two-way ANOVA with repeated measures was carried out to evaluate CBD’s modulation of cocaine seeking within the first session of extinction. We found that both, *CBD treatment* (F_1,65_ = 4.14; *p* < 0.05) and *time* (F_11,715_ = 7.65; *p* < 0.001) factors significantly influenced cocaine seeking during this specific session. Both factors significantly interacted (F_11,616_ = 2.36; *p* < 0.001), and a post-hoc analysis indicated that CBD effects became manifest after the first hour of session (VEH vs CBD, 70-min bin: Bonferroni; *p* < 0.01) (Figure 1d). ‘Extinction bursting’ behavior is considered as the paradoxical increase of operant responding when the instrumental behavior is no longer followed by the reinforcer presentation (Lerman et al., 2012; Skinner, 1938). To characterize this feature, we calculated a two-way repeated measures ANOVA for nosepoke responding during the FR1 phase, the first extinction session and the last extinction day. To better profile FR1 cocaine consumption, measures of this phase were defined as the mean number of active responses given in all sessions starting from the session in which acquisition criteria was reached. Results indicate that nosepoke responding changed across sessions (*session* factor: F_2,127_ = 89,97; *p* < 0.001) and was, overall, equivalent between the vehicle and CBD groups (*CBD treatment* factor: F_1,6_ = 3.22; *p* = 0.07). A significant interaction effect (F_2,127_ = 4,92; *p* < 0.01) evidences that CBD affected cocaine seeking depending on the session. Bonferroni’s post-hoc comparisons reveal that all groups exerted more nosepokes in the first extinction day than in the last FR1 session (Bonferroni; *p* < 0.001) and that the CBD group exhibited less cocaine seeking behavior in the first extinction day, compared to vehicle group (Bonferroni; *p* < 0.01) (Figure 1e). Furthermore, cocaine seeking behavior showed a significant decrease between the FR1 phase and the last extinction session in both groups (Bonferroni; *p* < 0.05), indicating that mice extinguished cocaine seeking behavior after undergoing the extinction protocol. Finally, our results show that the number of days required to reach extinction criteria did not differ among groups (unpaired *t* test: *t*_66_ = 0.01; *p* > 0.99) (Figure 1f).

**Figure 1.**
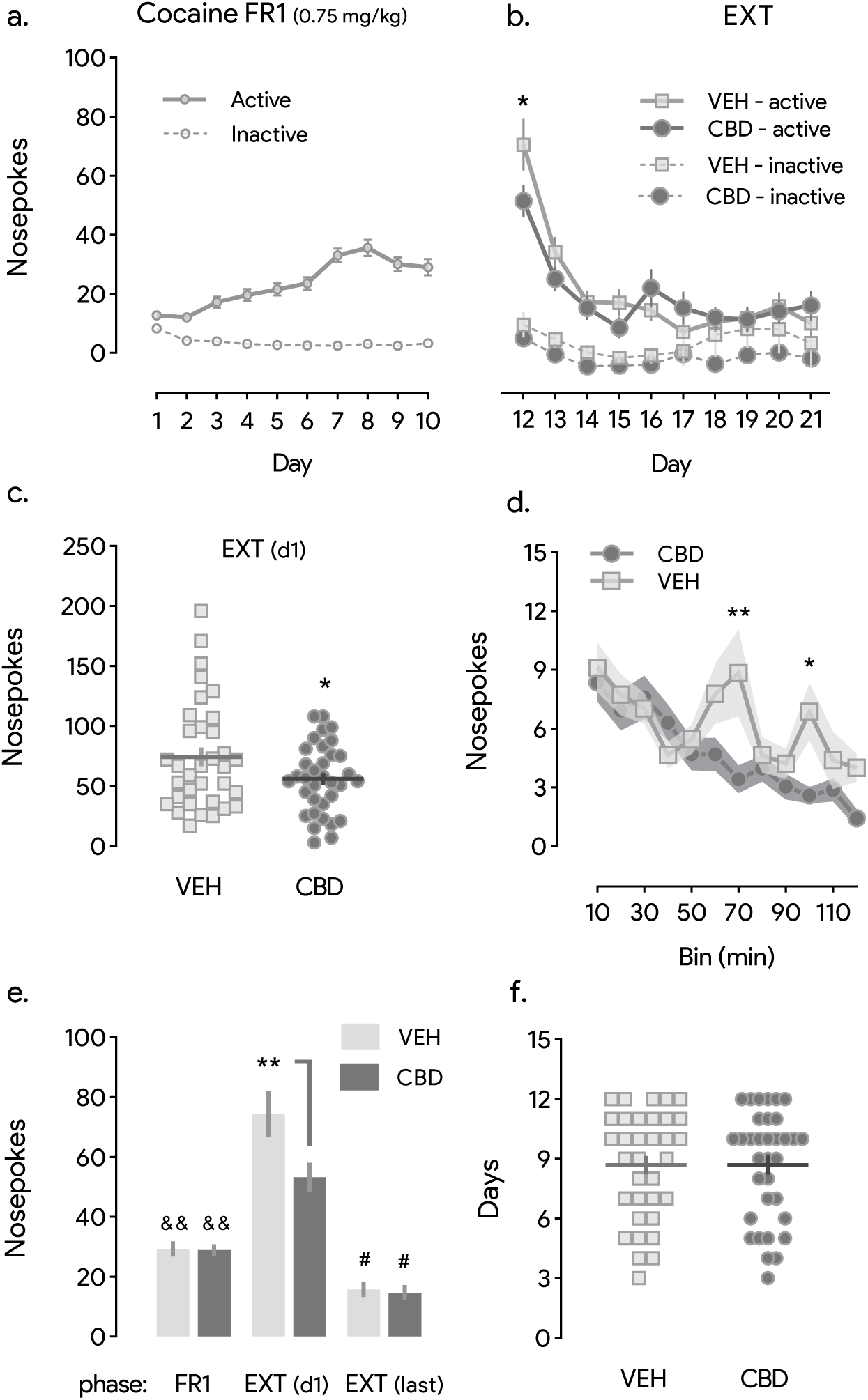
CBD specifically reduces ‘extinction burst’ responding, but not extinction learning development. **(a)** CD-1 male mice trained to seek the cocaine-associated nosepoke hole reached a stable pattern of self-administration after ten 2h-long sessions under a FR1 reinforcement schedule (n = 68). **(b)** CBD (20 mg/kg) treatment administered immediately before each extinction session reduces cocaine seeking behavior after the first administration (day 12 – active nosepokes, VEH vs CBD; Bonferroni, \**p* < 0.05) but does not modulate extinction learning progression (n = 34/group). **(c, d)** Cocaine seeking responding in extinction conditions is attenuated after a single CBD injection (20 mg/kg) (*t* test; \**p* < 0.05), an effect that becomes manifest following the first 70 minutes of session (same day, VEH vs CBD; Bonferroni, \*\**p* < 0.01, \**p* < 0.05) (n = 34/group). **(e)** ‘Extinction burst’ responding is observed in VEH- and CBD-treated groups, as cocaine seeking increases on the first extinction day in respect to the FR1 phase (Bonferroni, ^&&^*p* < 0.01). ‘Extinction burst’ responding is attenuated by CBD (20 mg/kg) (Bonferroni, \*\**p* < 0.01). Cocaine seeking responding declines after the extinction protocol, as evidenced by the difference observed between the FR1 phase and the last extinction day (Bonferroni, ^#^*p* < 0.05) (n = 34/group). **(f)** Despite differences in the first extinction session, the time required for each animal to reach extinction learning criteria is not affected by CBD (20 mg/kg) (n = 34/group). Data is represented as mean ± SEM. ‘EXT’: extinction. ‘EXT (d1)’: first day of extinction. ‘EXT (last)’: last days of extinction.

### CBD blocks cue-induced reinstatement of cocaine seeking behavior in a CB1-dependent manner

After reaching extinction criteria and performing a minimum of 8 sessions, 17 animals were randomly assigned to the cue-induced reinstatement condition. All mice underwent two consecutive reinstatement sessions separated 24h apart. They received AM4113 (5 mg/kg) in one session and vehicle on the other. The order of the injections was counterbalanced among subjects. As such, *AM4113 treatment* was considered a within-subject factor. Two determine the reinstatement of cocaine seeking behavior, a two-way ANOVA with repeated measures was calculated for nosepoke responding throughout the FR1 acquisition phase, the last extinction day and the session in which animals reinstated cocaine seeking behavior after receiving a vehicle injection. Analyses show a significant effect of the factors *CBD treatment* (F_1,15_ = 7.03; *p* = 0.181), *session* (F_2,30_ = 25.55; *p* < 0.001) and its interaction (F_2,30_ = 8.23; *p* < 0.01). The interaction effect was followed by Bonferroni’s post-hoc analyses revealing that CBD (20 mg/kg) specifically reduced cocaine seeking, compared to vehicle controls, in the cue-induced reinstatement session (Bonferroni; *p* < 0.001) (Figure 2a). Moreover, CBD-treated mice nosepoke responding in the reinstatement session did not significantly differ from the last extinction session (Bonferroni; *p* = 0.078). In contrast, vehicle mice significantly increased nosepoke responding in the reinstatement session, compared to the last extinction session (Bonferroni; *p* < 0.001). To determine if CBD attenuated cue-induced reinstatement of cocaine seeking in a CB1-dependent manner, we carried out a two-way, repeated measures ANOVA of seeking behavior throughout the two reinstatement sessions, one receiving vehicle and the other receiving AM4113 (5 mg/kg). *CBD treatment* was considered a between-subject factor and *AM4113 treatment* a within-subject factor. Results indicate that *AM4113 treatment* factor influenced nosepoking in reinstatement conditions (F_1,15_ = 7.24; *p* < 0.05). *CBD treatment* did not alter seeking behavior in all groups (F_1,15_ = 2.58; *p* < 0.5), but significantly interacted with *AM4113 treatment* (F_1, 15_ = 5.28; *p* < 0.05). Bonferroni’s post-hoc comparisons showed that CBD-treated mice exerted fewer nosepokes than vehicle-treated animals in the absence of AM4113 (Bonferroni; *p* < 0.05). However, when animals received AM4113, the difference between CBD and vehicle groups disappeared (Bonferroni; *p* > 0.999) (Figure 2b). AM4113 did not influence cocaine seeking behavior by itself, as vehicle-treated mice performed equally with or without AM4113 acute administration (Bonferroni; *p* > 0.999).

**Figure 2.**
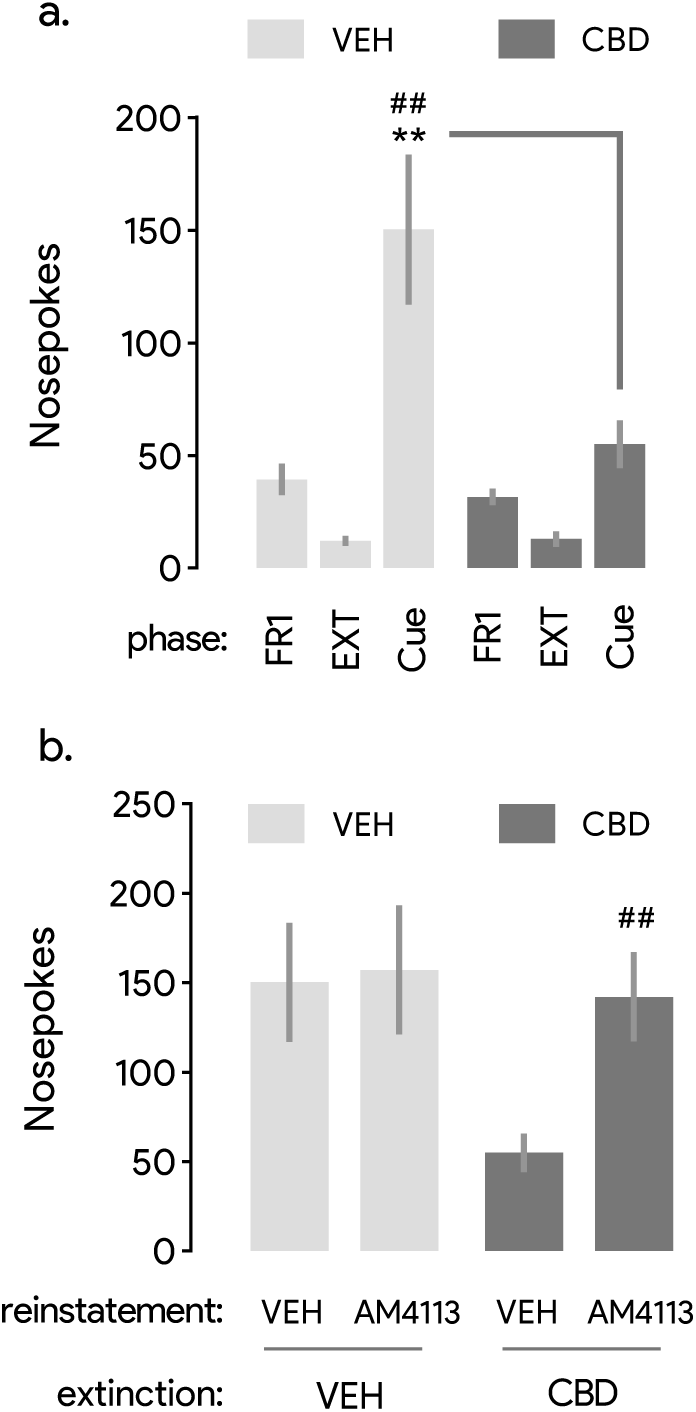
Cue-induced reinstatement is prevented in CBD-treated mice, an effect modulated by the pharmacological blockade of CB1 receptors. **(a)** Contingent cocaine-associated cue presentation did not induce reinstatement of previously extinguished seeking behavior in CBD-treated mice, contrary to the control group (FR1 phase vs cue-induced reinstatement session; Bonferroni, ^##^*p* < 0.01) (VEH, n = 9; CBD, n = 8). Cocaine seeking in the reinstatement session was reduced by extinction-targeted CBD (20 mg/kg) treatment (Bonferroni, \*\**p* < 0.01). **(b)** The CBD-induced reduction of cocaine seeking behavior in the cue-induced reinstatement test is reversed by the systemic administration of AM4113 (5 mg/kg) (CBD/VEH vs CBD/AM4113; Bonferroni, ^##^*p* < 0.01) (VEH, n = 9; CBD, n = 8). AM4113 administration does not influence nosepoking by itself. Data is represented as mean ± SEM. ‘EXT’: extinction.

### CBD facilitates the stress-induced reinstatement of cocaine seeking behavior in a CB1-dependent manner

After extinguishing cocaine seeking behavior, mice underwent two stress-induced reinstatement sessions in which they alternatively received AM4113 (5 mg/kg) or vehicle. In both sessions, cocaine seeking behavior was precipitated by the presentation of 5 mild footshocks (0.21 mA) within the operant chamber for 5 minutes. To characterize stress-induced reinstatement of cocaine seeking behavior in CBD-treated mice we ran a two-way ANOVA with repeated measures that yielded a significant participation of the *session* (F_2,18_ = 26.08; *p* < 0.001) and *CBD treatment* factors (F_1,9_ = 12.93; *p* < 0.01). A significant interaction between factors (F_2,18_ = 20.14; *p* < 0.001) allows to ascertain that only CBD-treated mice reinstated cocaine seeking behavior after footshock presentation (last extinction day vs reinstatement: Bonferroni; *p* < 0.001). Data suggests that control animals did not reinstated cocaine seeking behavior following footshock delivery (last extinction day vs reinstatement: Bonferroni; *p* = 0.677). Furthermore, the effects of CBD (20 mg/kg) on the reinstatement session were confirmed by comparing the magnitude of operant responding within this session between both groups (Bonferroni; *p* < 0.001) (Figure 3a). Moreover, the effects of the CB1 antagonist AM4113 (5 mg/kg) were evaluated using a counterbalanced, within-subject design along the two reinstatement sessions. A two-way ANOVA of nosepoke responding in these two sessions with *AM4113 treatment* as within-subject factor yielded a significant effect of *CBD treatment* (F_1,9_ = 23.39; *p* < 0.001), but not *AM4113 treatment* (F_1,9_ = 3.90; *p* = 0.07). However, as both factors interacted (F_1,9_ = 7.70; *p* < 0.05), Bonferroni’s post-hoc comparisons indicated that AM4113 (5 mg/kg) reversed the facilitating effects of CBD on stress-induced reinstatement (CBD/VEH vs CBD/AM4113: Bonferroni; *p* = 0.021) (Figure 3b). Conveniently, AM4113 (5 mg/kg) did not modulate cocaine seeking behavior reinstatement after footshock presentation by itself (VEH/VEH vs VEH/AM4113: Bonferroni; *p* > 0.999).

**Figure 3.**
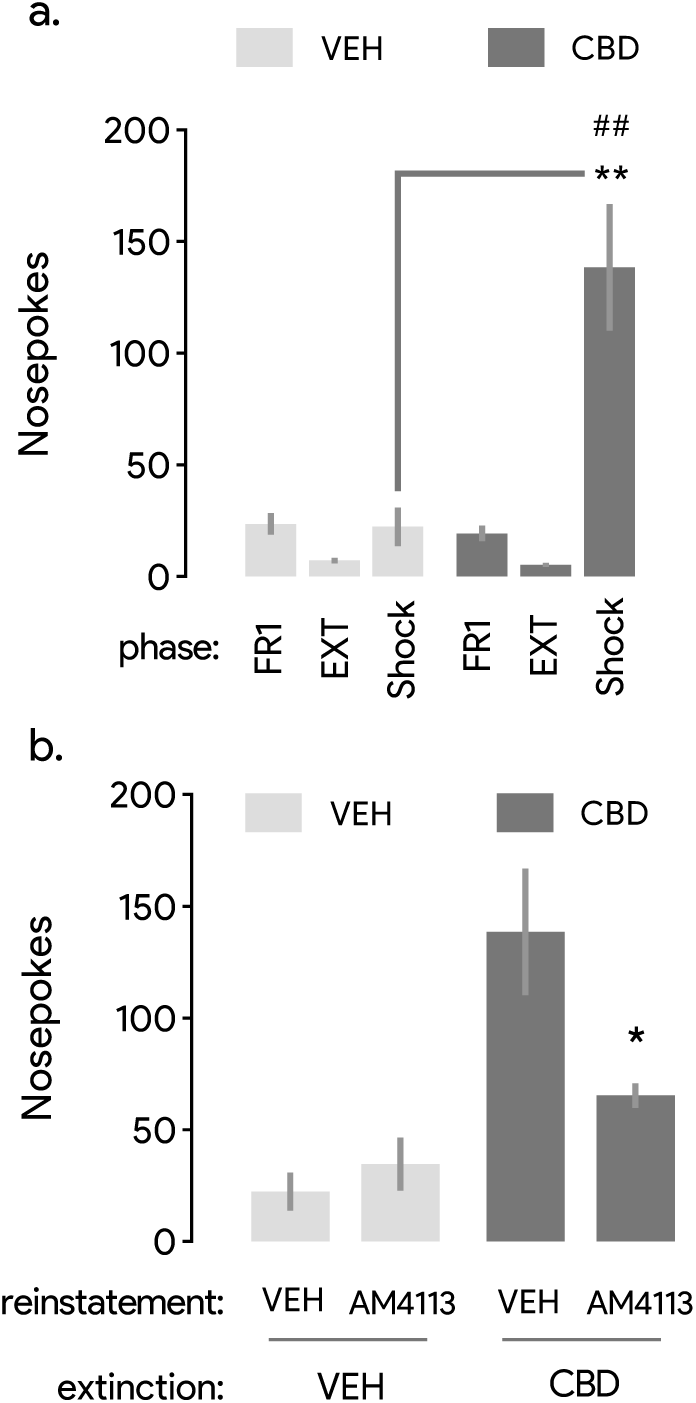
CBD extinction-targeted treatment facilitates stress-induced reinstatement of cocaine seeking in a CB1-dependent manner. **(a)** Compared to the control group, animals receiving CBD (20 mg/kg) during the extinction phase show increased expression of cocaine seeking behavior after repeated footshock (0.21 mA) presentation (Bonferroni, \*\**p* < 0.01). Compared to the last extinction day, nosepoke measures in the reinstatement test increased in the CBD group (Bonferroni, ^##^*p* < 0.01), but not in the VEH group (VEH, n = 6; CBD, n = 5). **(b)** The facilitating effect of CBD (20 mg/kg) on stress-induced reinstatement is reduced by the systemic administration of AM4113 (Bonferroni, \**p* < 0.05) (VEH, n = 6; CBD, n = 5). AM4113 administration does not influence nosepoking by itself. Data is represented as mean ± SEM. ‘EXT’: extinction.

### CBD does not influence cocaine-primed reinstatement of cocaine seeking behavior

After extinction, cocaine seeking behavior was reinstated by administering an i.p. injection of cocaine (10 mg/kg) immediately before each reinstatement session. A two-way, repeated measures ANOVA of nosepoke responding along the FR1 phase, last extinction session and reinstatement session without AM4113 was calculated in order to elucidate whether animals recovered initial levels of cocaine seeking after drug priming. A significant effect of the *session* factor (F_2,30_ = 7.52; *p* < 0.05) indicated that cocaine responding varied across experimental phases (Figure 4a). However, the *CBD treatment* (F_1,15_ = 1.15; *p* = 0.30), and the interaction between factors (F_2,30_ = 0.58; *p* = 0.561) did not show significant effects. Nonetheless, the *session* factor effect was significant, and thus it allowed us to perform a session-specific post-hoc test for all animals, independently from the group. Results indicated that, overall, nosepoke responding was higher in the reinstatement session than in the last extinction session, indicating that drug-induced reinstatement occurred in both groups. Figure 4b shows the absence of AM4113 (5 mg/kg) effects on the reinstatement of cocaine seeking behavior after drug priming. A two-way, repeated measures ANOVA disclosed *CBD treatment* (F_1,15_ = 0.26; *p* = 0.614) and *AM4113 treatment* (F_1,15_ = 1.83; *p* = 0.196) as non-significant factors. There was no interaction between main factors (F_1,15_ = 3.95; *p* = 0.065).

**Figure 4.**
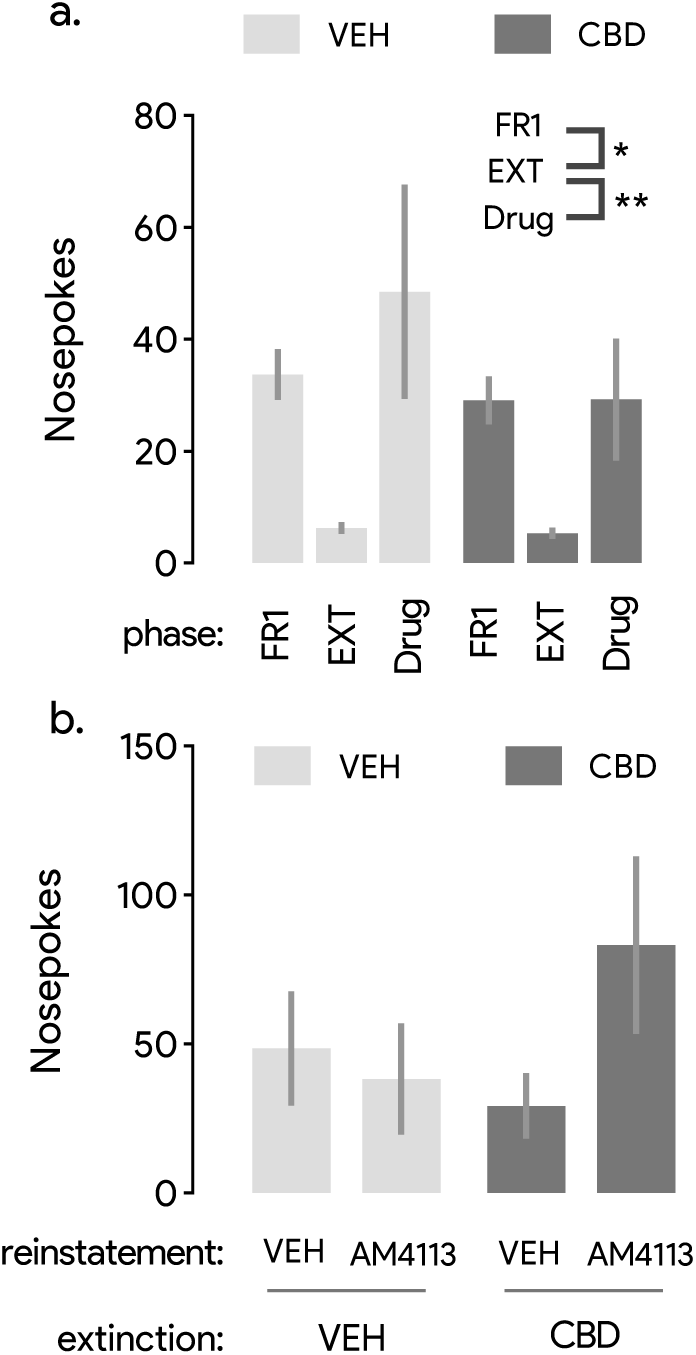
CBD treatment has no effects on the drug-induced reinstatement of cocaine seeking behavior. **(a)** VEH and CBD groups equally reinstate cocaine seeking responding precipitated by a cocaine priming (i.p., 10 mg/kg) (group-unspecific, FR1 phase vs drug-induced reinstatement; Bonferroni, \*\**p* < 0.01, \**p* < 0.05) (VEH, n = 8; CBD, n = 9). **(b)** The acute, systemic administration of the CB1 antagonist AM4113 has no effects on cocaine seeking behavior in a drug-induced reinstatement test (VEH, n = 8; CBD, n = 9). Data is represented as mean ± SEM. ‘EXT’: extinction.

## DISCUSSION

The results obtained herein document an unexpected differential modulation of the reinstatement of cocaine seeking, following CBD (20 mg/kg) treatment during extinction learning. Extinction-targeted CBD treatment did not drastically affect extinction learning but decreased ‘extinction burst’ responding and cue-induced reinstatement of cocaine seeking. Conversely, the same CBD treatment facilitated cocaine seeking precipitated by a footshock presentation. Finally, CBD did not influence cocaine-seeking behavior reinstated after re-exposure to a priming dose of the drug. Furthermore, results suggest a participation of CB1 receptors in the behavioral manifestation of CBD-induced neuroplasticity, as measured in the cocaine seeking reinstatement tests. Therefore, CB1 receptor antagonist AM4113 (5 mg/kg) blunted the facilitation and decrease of cocaine seeking behavior in the stress- and cue-induced reinstatement phases, respectively.

We treated cocaine self-administering mice with CBD (20 mg/kg) during an extinction-based period of abstinence to the drug. Extinction of operant drug seeking behavior is the most used preclinical paradigm to model drug use cessation (Bossert et al., 2013). For this reason, manipulations that strengthen extinction memories have been proposed as useful strategies that help to maintain abstinence (Everitt, 2014; Farrell et al., 2018; Kalivas and Volkow, 2011). Our results show a lack of effect of CBD (20 mg/kg) on cocaine seeking behaviors during extinction conditions. Studies assessing the potential protective effects of CBD in psychostimulant-consuming rodents are still scarce. For this reason, comparisons with other previous works are limited. The closest approximate was reported in Luján et al. (2018). In this previous study, we showed that animals treated with CBD (20 mg/kg) during the acquisition phase of cocaine operant taking extinguished seeking behavior in a similar extent than control mice. Similarly, CBD (60 mg/kg) did not facilitate extinction learning of ethanol taking behavior in rats (Viudez-Martínez et al., 2018). Another study using heroin also described that CBD (10, 20 mg/kg) did not affect lever pressing during extinction training (Ren et al., 2009). Altogether, results do not support CBD as a potential therapeutic solution to promote extinction-based abstinence in rodent models of drug abuse. Therefore, CBD would not be exerting its putative protective effects against maladaptive behavioral manifestations of cocaine reinforcement by facilitating the development of extinction memories.

Despite an overall lack of effect of CBD on extinction training, results show a decrease in cocaine seeking behavior after one CBD (20 mg/kg) injection, in the first extinction day. In this session, experimental subjects typically show an ‘extinction burst’ in which operant responses are greatly increased in respect to the last session when cocaine was still available (Lerman et al., 2012; Skinner, 1938). Such ‘extinction bursting’ behavior is often considered as a measure of the urge to consume the drug (craving) that is not affected by the acute pharmacological effects of the drug or the development of a previous extinction learning and thus, is viewed as a predictor of relapse behavior (Wee et al., 2012). Such an effect was not reported in the previous studies evaluating the effects of CBD treatment on extinction learning (Ren et al., 2009; Viudez-Martínez et al., 2018). Similarly, Mahmud et al. (2017) did not find that CBD (5, 10 mg/kg) reduced cocaine seeking in the first withdrawal session. However, transdermal CBD (15 mg/kg) induced an effect similar to what we report (Gonzalez-Cuevas et al., 2018). After the first exposure to CBD, cocaine seeking was found decreased. In drug self-administration regimes in which cocaine is infused at very low doses (0.03 mg/kg) or is scarcely administered (as in a progressive ratio reinforcement schedule), an acute injection of CBD (20, 40 mg/kg) induced the most effective attenuating effects on operant responding (Galaj et al., 2019). CBD’s protective effects centered at the initial stages of cocaine abstinence could represent a beneficial feature that should warrant further experimental exploration. Indeed, clinical trials indicate that behavioral measures of drug craving during early abstinence negatively correlate with treatment retention and outcome (Ahmadi et al., 2009; Kampman et al., 2001; Patkar et al., 2002; Poling et al., 2007). However, there are no experimental medications being currently tested in humans that show desirable effects in the early stages of abstinence (Kampman, 2019).

In humans, places and stimuli that were experienced during drug use trigger feelings of craving and therefore, pose a major risk factor contributing to relapse. In rodents, associated cues can be conditioned to the delivery of the drug. Hence, after extinguishing drug seeking, re-exposure to paired drugs can elicit reinstatement of the seeking behavior. In our study, we evaluated the effects of CBD (20 mg/kg) in a mouse model that aims to recapitulate the relapse-promoting properties of paired cues after a period of extinction-based abstinence (Venniro et al., 2016). Results indicate that CBD (20 mg/kg) specifically blocks cue-induced cocaine seeking reinstatement. Such a protective role of CBD in similar experimental conditions has been previously reported using rats. A transdermal preparation of CBD (15 mg/kg, 7 days) implanted after the first reinstatement of cocaine seeking attenuated subsequent measures of motivation to pursue cocaine, and alcohol, when related cues (context) were presented (Gonzalez-Cuevas et al., 2018). In a non-operant model of cocaine reinforcement, CBD was also reported able to impair the reconsolidation of cocaine-induced conditioning place preference (de Carvalho and Takahashi, 2016). In conclusion, the results here presented expand upon the available evidence suggesting a protective role of CBD against vulnerability states promoting relapse in conditions associated to re-exposure of cocaine-associated cues.

As cues, stressful events are also associated with higher risk of relapse in abstinent individuals (Koob and Le Moal, 2005). In rodents, acute stressors such as footshock presentation or yohimbine injections can reinstate a previously extinguished cocaine seeking behavior (Mantsch et al., 2016). Here, we found that CBD (20 mg/kg) extinction-targeted treatment unexpectedly facilitated stress-induced reinstatement of cocaine seeking behavior. Such an adverse consequence of CBD has not been reported before. Instead, Gonzalez-Cuevas et al. (2018) reported that CBD (15 mg/kg) attenuated cocaine pursue after yohimbine priming. Yohimbine (a pharmacological alpha 2 blocker that increases peripheral and central effects of stress) is considered equivalent to stress (Bossert et al., 2013), but evidence suggests that its effects are not the same that those induced by a footshock presentation (Chen et al., 2015). Thus, such protocol difference could explain divergencies between results. Additionally, the parameters of both CBD treatments considerably differed. While we administered CBD by i.p. route, Gonzalez-Cuevas et al. (2018) implanted it as a transdermal application. Moreover, while we tested the effect of an extinction-targeted CBD treatment, theirs study was focused on the effects of CBD after cocaine seeking behavior was reinstated. Hence, this opens the possibility that CBD-induced neuroplasticity might interfere with extinction learning neurocircuitry in a way that could facilitate the expression of cocaine memories in stressful conditions. Another possibility is that CBD-treated mice performed the stress-induced reinstatement test in an elevated state of anxiety. Although CBD shows prominent anxiolytic effects (Gonzalez-Cuevas et al., 2018; Luján et al., 2019, 2018), a recent report indicated that prolonged CBD treatments could induce anxiety-like states (Schleicher et al., 2019). Further experimental efforts will be needed to test this specific hypothesis. Even though, our results raise the issue of CBD extinction-based treatment being counterproductive for relapse prevention under certain circumstances. In case of being confirmed, such an effect could importantly limit the therapeutical applications of CBD in cocaine addiction.

A low priming dose of cocaine in an abstinent subject can elicit drug craving and contribute to increased risk of relapse. Priming doses of cocaine (< 15 mg/kg) can also reinstate previously extinguished cocaine seeking behavior in rodents (Shaham et al., 2003). Here, we evaluated the reinstatement-promoting effects of a priming dose of cocaine (10 mg/kg) in animals that received a repeated CBD treatment (20 mg/kg) during extinction. Results indicate that CBD did not influence the drug-induced reinstatement of cocaine seeking behavior in mice. Previous reports showed that different treatments of CBD could attenuate this response in rats. In the study of Gonzalez-Cuevas et al. (2018), a transdermal preparation of CBD (15 mg/kg, 7 days) reduced cocaine-induced drug seeking reinstatement in rats. Similarly, primming-induced reinstatement of methamphetamine seeking was found attenuated following acute intracerebroventricular CBD (µg/5µl) (Karimi-Haghighi and Haghparast, 2018) or acute i.p. CBD (80 mg/kg) administration. In the latter study, lower doses of CBD (20, 40 mg/kg) did not reduce operant responding. We have previously shown that primming-induced reinstatement of cocaine seeking behavior was not affected in mice treated with CBD (20 mg/kg, 10 days) during the acquisition phase of self-administration behavior (Luján et al., 2018). In a different paradigm, acute CBD (5 mg/kg) did not prevent the reward-facilitating effects of cocaine in an intra-cranial self-stimulation study (Katsidoni et al., 2013). Also, Ren et al. (2009) found that CBD (10, 20 mg/kg, 5 days) was unable to reduce drug-induced reinstatement of heroin seeking behavior. So far, data is still inconclusive to confirm the therapeutic potential of CBD to reduce drug-primed relapse in rodent models, especially considering the divergences observed between species and drugs.

Cocaine-evoked plasticity in the mesocorticolimbic system, particularly within the nucleus accumbens (Nac), drives drug-adaptive behavior (Nestler and Lüscher, 2019; Pascoli et al., 2015). CBD is thought to interact with cocaine-induced neuroplasticity in multiple ways (Calpe-López et al., 2019). However, no study has yet directly assessed the participation of a putative CBD mechanism in a rodent model of drug operant seeking in relapse conditions. To identify such a mechanism, we explored the role of CB1 receptor signaling in the behavioral expression of cocaine memories in CBD-exposed mice. Commonly used CB1 antagonists often decrease the reinforcing properties of drugs of abuse by themselves (Bi et al., 2019). To avoid this, we used the neutral antagonist AM4113 (5 mg/kg) at a dose that has been probed to decrease CB1 receptor activity without inducing inverse agonist activity (He et al., 2018). Hence, in the reinstatement tests here employed, AM4113 did not exert any effects in control animals. This was also observed in the CBD group that showed no differences with control animals on drug-induced cocaine seeking reinstatement.

The attenuation of cue-induced reinstatement by CBD (20 mg/kg) was blocked with the CB1 receptor antagonist AM4113 (5 mg/kg). Other works hinted at a possible participation of CB1 receptors in the protective effects of CBD in drug self-administration studies. Striatal CB1 mRNA expression was found reduced following a CBD treatment with drug seeking attenuating effects (Ren et al., 2009; Viudez-Martínez et al., 2018). The effects of AM4113 on CBD-induced changes during extinction learning indicates that the expression of these effects are modulated by CB1 cannabinoids receptors on reinstatement conditions. A possible interpretation of these effects arises considering the ability of CBD to normalize the alterations induced by drugs of abuse in the glutamatergic system. Accompanying the CB1 receptor changes, CBD also influences α-amino-3-hydroxy-5-methyl-4-isoxazolepropionic acid receptor (AMPAR) subunit composition (Luján et al., 2018; Ren et al., 2009), reducing the amount of calcium-permeable AMPAR (CP-AMPAR). CP-AMPAR accumulation occurs during cocaine withdrawal and contributes to craving incubation (Ma et al., 2014). Crucially, the expression of CP-AMPAR-dependent alterations in reinstatement conditions is regulated by CB1 receptors and the metabotropic glutamate receptor 1 (mGluR5) (Wolf, 2016). In particular, mGluR5 and CB1 receptors act as a negative feedback of the increased excitatory output produced by CP-AMPAR accumulation in Nac synapses (Katona and Freund, 2008; Loweth et al., 2014). Therefore, if CBD attenuates cue-induced reinstatement of cocaine seeking by preventing the accumulation of CP-AMPAR during withdrawal, it is plausible that this effect disappears when the remaining CP-AMPAR can be increasingly activated as a consequence of CB1 receptor blockade by AM4113. If that were to occur, the expression of drug craving, here elicited by cue presentation, will increase reaching control levels (as here reported).

Similarly, our results suggest that CB1 receptor activity is also crucial to the expression of CBD’s facilitation of stress-induced reinstatement of cocaine seeking. Given that CBD shows important molecular and behavioral anti-anxiety effects (Campos et al., 2017; Skelley et al., 2019), it is difficult to propose a possible mechanism underlying these results. However, the structural and functional ubiquity of CB1 receptors in the central nervous system (Busquets-Garcia et al., 2018) allows CBD to interfere in unexpected ways with different brain structures, in certain environmental situations leading to drug relapse. In this sense, it is possible that the diverse effects of CBD over CB1-expressing, heterogeneous neuronal populations may lead to both undesirable and protective consequences. CBD can have opposed effects on CB1 receptors, given that it can act as an indirect agonist (De Petrocellis et al., 2011) and as a biased negative allosteric modulator (Tham et al., 2019). The complex pharmacology of this phytocannabinoid possess interesting properties for the management of different neuropsychiatric conditions but can also lead to some undesirable effects that are especially hard to predict (Brown and Winterstein, 2019). In conclusion, the complex pharmacology of CBD over CB1 receptor is still poorly understood and, as we show here, unexpected consequences can arise in specific situations leading to cocaine seeking reinstatement in rodent models of operant drug seeking. These effects should be considered in the risk-benefit assessment of CBD as a potential pharmacotherapy for the treatment of cocaine use disorders.

**ABBREVIATIONS**
5-HT_1A_: 5-hydroxytryptamine 1 A receptor
AEA: anandamide
AMPA: α-amino-3-hydroxy-5-methyl-4-isoxazolepropionic acid receptor
CB1: cannabinoid receptor type 1
CB2: cannabinoid receptor type 2
CBD: cannabidiol
FAAH: fatty acid amid hydrolase
FR1: fixed ratio 1
GPR55: G-protein receptor 55
mGluR5: metabotropic glutamate receptor 5
Nac: nucleus accumbens
PPARγ: peroxisome proliferator-activated gamma
TRPV1: transient receptor potential vanilloid 1 receptor

## ACKNOWLEDGEMENTS

This work was supported by Ministerio de Economía y Competitividad (grant number SAF2016-75966-R-FEDER), Ministerio de Sanidad, Asuntos Sociales e igualdad (Retic-ISCIII-RD/16/0017/0010-FEDER and Plan Nacional Sobre Drogas (#2018/007), and by. M.A.L. received FPU grant (15/02492) from the Ministerio de Educación, Cultura y Deporte. L.A.-Z. received FPI grant (BES-2017-080066) from the Ministerio de Economía y Competitividad.

The authors declare no conflict of interest.

